# An Experimental Validation of the Computer Fluid Dynamics of Normal Human Nasal Airflow Using a Precise 3-D Printing Model

**DOI:** 10.1101/2020.03.19.998757

**Authors:** Xiaoli Fu, Yin Cheng, Huanhai Liu, Jianchun Liao, Mingqing Kou, Chao Shen, Qinghua Ge, Shenglin Yan, Hu Peng

## Abstract

Computer Fluid Dynamics (CFD) is a popular method for studying airflow of nasal cavities. However, the data of CFD studies has rarely been validated through experiments. To test the accuracy of CFD computation, we studied the consistency of the air pressure of nasal cavities in the CFD and the experiment. A proportional resin model of a normal human subject’s nasal cavities was created by a 3-d printer with a precision value of 0.1mm. The pressure of 63 check points in the nasal cavities in different breathing states was measured. The experimental data was compared with the data obtained by CFD simulation. At the flow rates of 180 ml s^-1^ and 560 ml s^-1^, the pressure in all check points remained highly consistent with the CFD data. At 1100 ml s^-1^ flow rate, there was a significant deviation in the posterior segment of the nasal cavity during exhalation. The method used in this study to measure the pressure in the nasal cavities can be used in experimental validation of CFD data. The computational methods and the boundary conditions used in this study resulted in a high agreement between the results of the CFD simulation and the experiment.

**Author Summary:** In the contemporary era, Computer fluid dynamics (CFD) is the mainstream method for studying air flow. Due to the complex anatomical structure of the nasal cavity, the CFD results of the nasal flow have rarely been experimentally verified. This study provides a method to verify the methods and results of nasal CFD. We printed an accurate model of a normal person’s nasal cavity with a high-precision 3D printer. In this nasal cavity model, we set 63 small holes to detect the air pressure of the places we concerned. Three different nasal flow quantity are used to represent different breathing conditions: high (1100 ml s-1), medium (560 ml s-1), and low (180 ml s-1). In medium and low nasal flow quantities, our CFD results are in good agreement with the experimental pressure values. On this basis, we analyzed the characteristics of nasal airflow in normal people. The method used in this study to measure the pressure in the nasal cavities can be used in experimental measurements of the partial resistance of the nasal cavity. With proper modification, it can be applied to the clinical practice for nasal resistance, giving more help for the design of the operation plan.

## Introduction

As the pathway of the respiratory system, the nasal cavity has important physiological functions of passing air, filtering air, regulating temperature and humidity of air, sensing smell, and providing immune defense. The change in the anatomic structure of the nasal cavity leads to the change in the airflow of the nasal cavity. This can cause the change of the physiological functions of the paranasal sinuses or even illnesses. The study of the internal airflow in the nasal cavity is very important in terms of guiding surgeries[1]. As far as the current clinical practice is concerned, only limited measurements of the nasal flow parameters (such as rhinomanometry, acoustic rhinometry) can be taken, and correlation between these measurements and clinical practice is questionable[2].

Due to the complex internal structure of the nasal cavity, it is difficulty to study its internal flow directly. The internal flow in the nasal cavity can only be estimated indirectly by studying the flow outside the nasal cavity[3]. Previous researchers Hahn et al. made a valuable study by constructing an enlarged scale (20x) model of a human nasal cavity[4]. Another method used was to study the nasal flow through cadavers[5]. Resin model was one of the models employed in previous studies[6]. With the development of computer technology, more researchers were using finite element methods to create computer models and the CFD method to simulate the airflow in the nasal cavity[6-8]. Not limited by the anatomical inaccessibility of the nasal cavity, CFD enables the calculation of various parameters of the intranasal flow field with a high temporal and spatial resolution[2]. This means computer modeling can provide detailed and objective characteristics of airflow for all locations and periods of an individual nose. Furthermore, the ability to remodel computer simulations provides a potential predictive tool for planning nasal surgeries[9]. As the methods used by researchers are not entirely the same, under what circumstance a method is more suitable for nasal cavity studies needs further investigations[2, 10-15]. Although there have been some studies comparing CFD simulation and experimental data[4, 6, 16, 17], the models used are not accurate enough. Therefore, they cannot fully explain the pros and cons among the existing computation methods.

An adult’s inspiratory nasal flow rate in restful breathing state is within the range of 5-12L min^-1^[18]. To find out whether the breathing equations for high (1100 ml s-1), medium (560 ml s^-1^), and low (180 ml s^-1^) nasal flow rates are accurate, we made an experimental measurement of a normal person’s nasal cavity model, which was 3-d printed with 1 : 1 precision ratio, and compared the results with the data obtained through CFD simulation. ml s^-1^

## Results

### 1. Normal nasal cavity model of CFD and 3-D printing

We first built a model of nasal cavity from a healthy female Chinese (35 years old). The model in IGES format is shown in Fig 1A. The three blue lines were connected by the air pressure check points which were located in superior, middle, inferior nasal meatuses. The details of the grid for the nasal cavity surface are shown in Fig 1B. The coordinate system of the model is showed in Fig 1C. The 1:1 3D printing model, with holes inside for air pressure check, are shown in Fig 1D(a). The experimental device is shown in Fig 1D(b).

**Fig 1.**
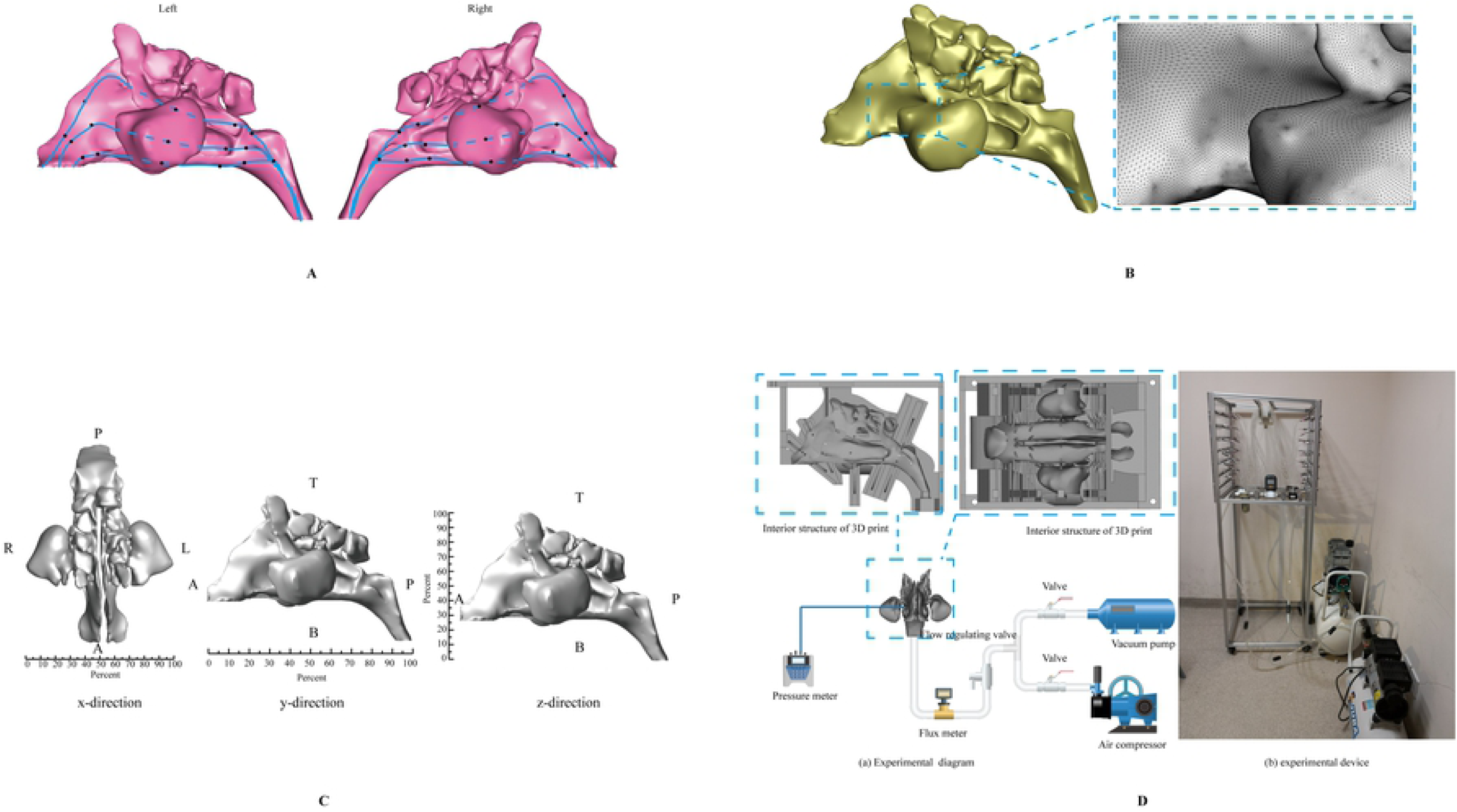
Nasal model and experiment equipment. A. Nasal cavity model of a Chinese health female. B. The grid for the nasal cavity surface. C. Coordinate system of the nasal model. D (a) Experimental flow chart, (b) Experimental device.

### 2. Air pressures in most positions of nasal cavity were consistent in CFD and experiment

The data of air pressure in the points of superior, middle, inferior nasal meatus simulated by CFD and checked by experiment are shown in Fig 2A and Fig 2B. The experimental data and the CFD data maintained a good consistency at the pressure of each test point at the flow rates of 180 ml s^-1^ and 560 ml s^-1^. At the flow rate of 1100 ml s^-1^, the experimental data and the simulated data were basically the same, and there was a deviation in the posterior segment of the nasal cavity during exhalation.

**Fig 2.**
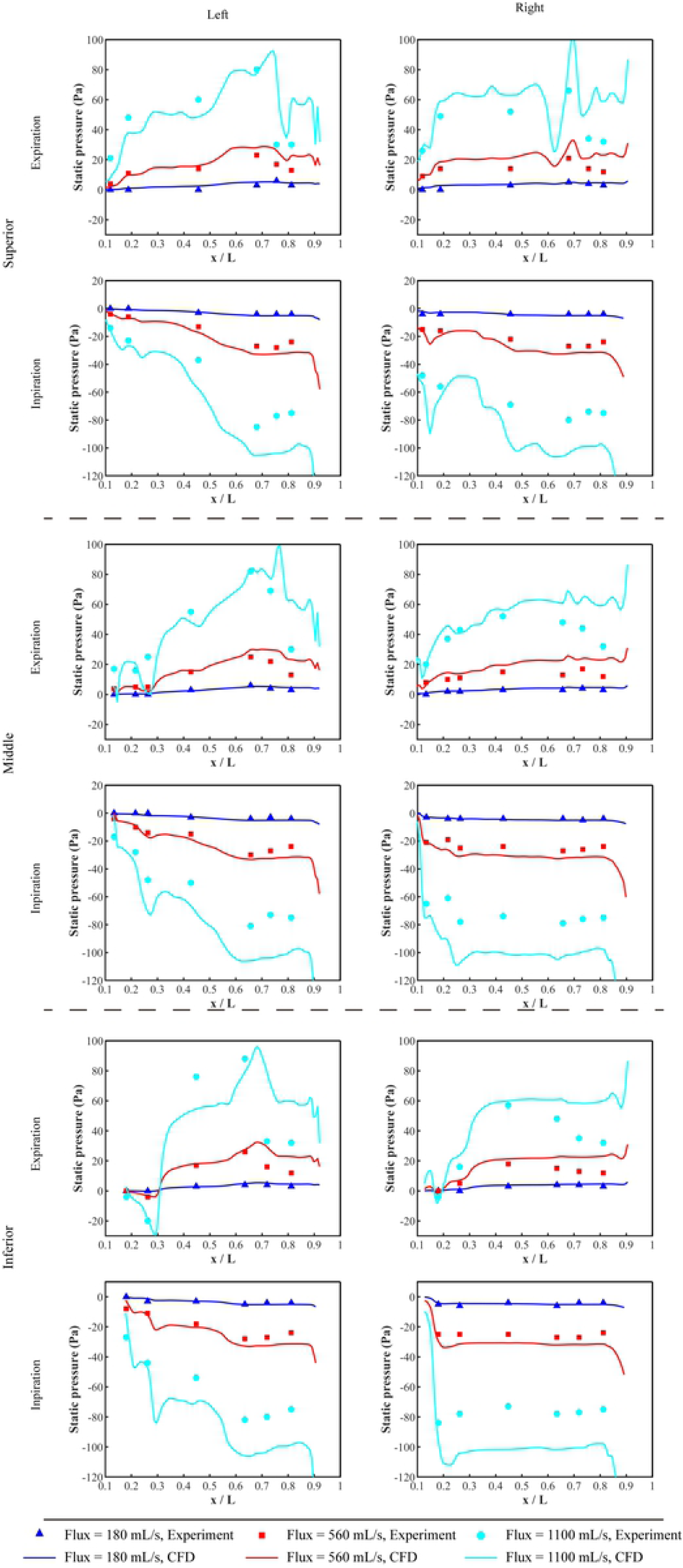
Experimental and CFD data of static pressure in nasal cavity

Experimental and CFD data of static pressure in the nasal cavity during exhalation and inhalation at different flow rates (180 ml s^-1^, 560 ml s^-1^ and 1100 ml s^-1^).As shown in Fig 2, the simulation data was close to the experimental results. When the flow rate was 180 ml s^-1^, the error was the smallest. With the increase of the flow rate, the error increased gradually. With the increase of the flow rate, the pressure fluctuated greatly. When exhaling, the area with the highest pressure was located in the range of X / L = 0.6 ∼ 0.8 of the middle nasal meatus. When inhaling, the area with the lowest pressure was in the range of X / L = 0.6 ∼ 0.7 of the middle meatus.

There was a slight difference between the left and right nasal meatuses. In the superior nasal meatus, the pressure on the left side was slightly lower than that on the right side when exhaling. When inhaling, the pressure difference on the left and right sides was small at the flow rate of 180 ml s^-1^ or 560 ml s^-1^. However, at the flow rate of 1100 ml s^-1^, the pressure fluctuation in the right nasal meatus was much larger. In the middle nasal meatus, the pressure of the left side was slightly higher than that on the right side when exhaling, and the pressure difference on the left and right sides was smaller when inhaling. In the inferior nasal meatus, the pressure on the left side was slightly higher than that on the right side when exhaling, whereas the pressure difference between the left and right sides was small at the flow rate of 180 ml s^-1^ or 560 ml s^-1^ when inhaling. However, at the flow rate of 1100 ml s^-1^, the pressure fluctuation of the left inferior nasal meatus was larger.

### 3. Distributions of the relative pressure during inspiration and expiration

The pressure distribution at the flow rates of 180, 560 and 1100 ml s^-1^ is shown in Fig 3A-C. When exhaling, the internal pressure of the nasal cavity was positive. The pressure increased gradually from the outside of the nostril to the inside of the nasal cavity. In contrast, the pressure was negative when inhaling.

**Fig 3.**
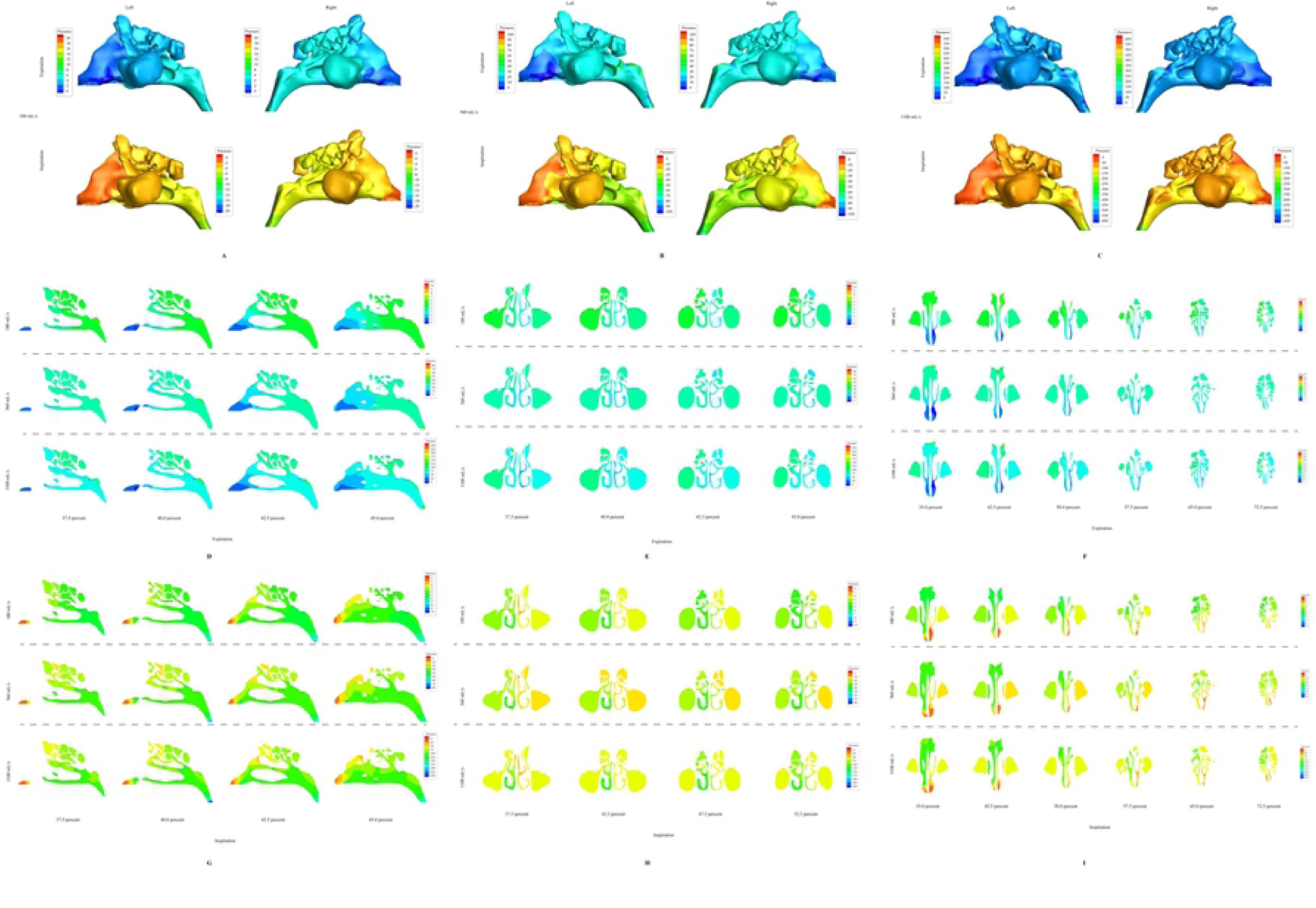
Stress nephogram.of nasal flow. A-C. Stress nephogram of exhaling and inhaling at different flow rates (180 ml s^-1^, 560 ml s^-1^ and 1100 ml s^-1^). D-F. Stress nephogram on different sections of X, Y and Z axes during exhalation. G-I Stress nephogram on different sections of X, Y and Z axes during inspiration.

As observed from the X-axis direction (Fig 3D and G), under the condition of small flow rate (180 ml s^-1^), the pressure changes at exhalation and inhalation mainly occur in the anterior part of nasal cavities. With the increase of the flow rate (560 ml s^-1^ and 1100 ml s^-1^), the area of pressure changes gradually expand to the whole nasal meatuses.

As observed from the Y-axis and Z-axis direction (Fig 3E,F,H and I), under the flow rate of 180 ml s^-1^, the pressure difference between the left and right nasal passages was smallest when exhaling. With the increase of the flow rate, the pressure difference between the left and right nasal meatuses increased gradually. However, whether the flow rate was small or large, the pressure difference between the left and right nasal meatuses was larger in inhaling than that in exhaling. This difference can be seen more clearly from the direction.

### 4. Distributions of the flow velocity

In the velocity nephogram of inhalation (Fig 4A), as the flow rate increased, the flow velocity also continuously increased. But the maximum velocity of inhalation occured in the front of inferior nasal meatus at the three flow rates. It is obvious that the flow velocity in the right nasal cavity is larger than that in the left.

**Fig 4.**
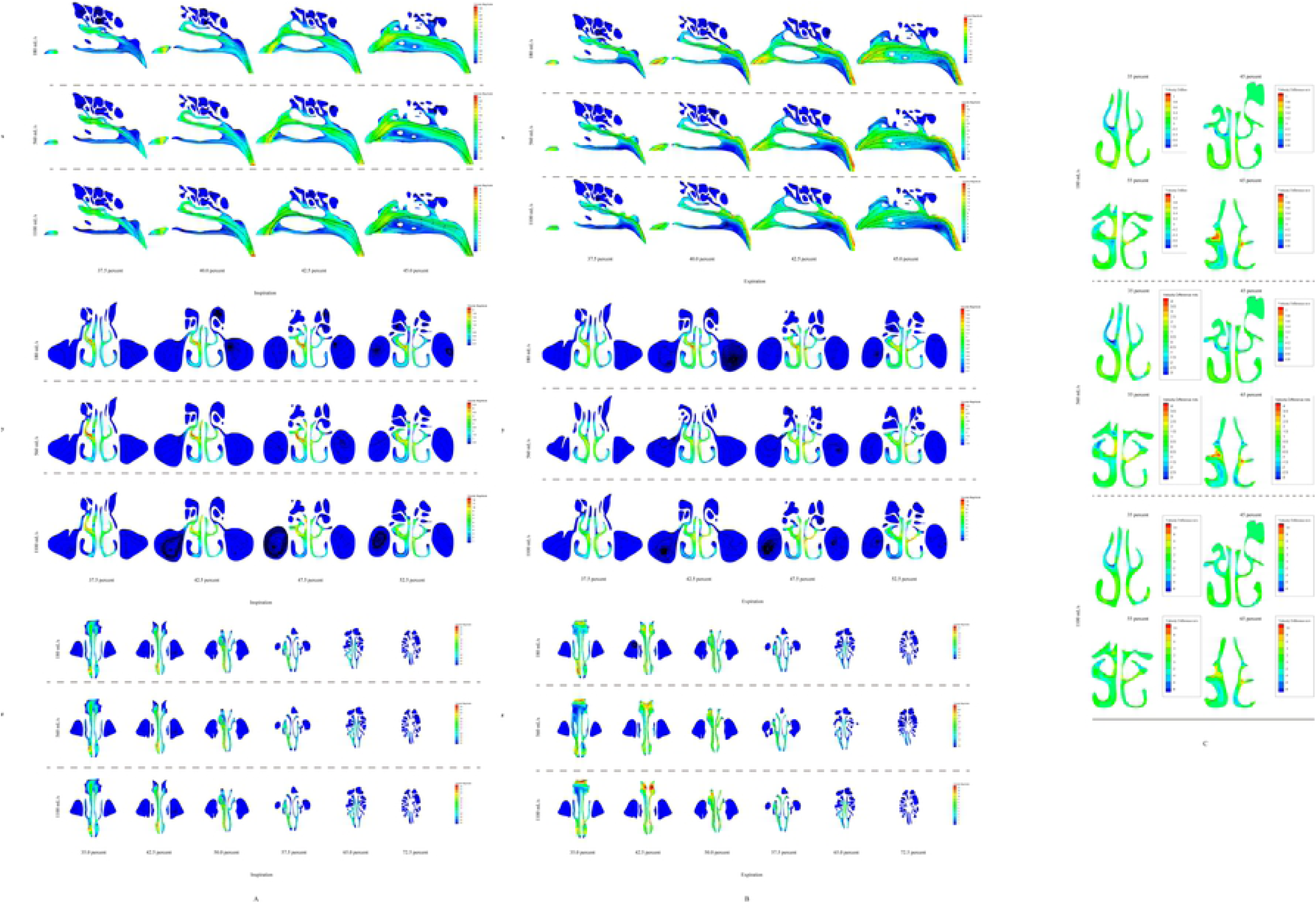
Distributions of the flow velocity in nasal cavity. A. Inspiration flow velocity on different sections of X, Y and Z axes. B. Expiration flow velocity on different sections of X, Y and Z axes. C. The difference value of flow velocity between inspiration and expiration in the same position.

The velocity nephogram of exhalation (Fig 4B) indicates that the maximum velocity of exhalation also occured in the front of inferior middle nasal meatus. From the Y-axis direction, it shows clearly that the flow velocity in the right nasal cavity was greater than that in the left nasal cavity under the flow rate of 180 ml s^-1^ when exhaling, which was consistent with inhalation. However, as the flow rate growing, the flow velocity in left middle nasal meatus inreased much more rapid than that the right.

Fig 4C shows the distributions of velocity difference. The velocity difference is the absolute value of expiratory velocity minus the absolute value of inspiratory velocity at the same position. At the cross sections of 35% and 45%, the difference in respiratory velocity near the middle turbinate was the largest, and it is consistent for all three flow rates, meaning that the expiratory velocity is significantly lower than the inspiratory velocity here.

### 5. Distributions of the nasal flow

Fig 5A shows the division of the superior, middle, inferior nasal meatuses in this article on the coronal section of the nasal cavity at different positions. The area of the left and right nasal meatuses at different slices are shown in Table 2.

**Table 1.** statistics of slice areas

| Slice location | Left |  |  |  | Right |  |  |  |
| --- | --- | --- | --- | --- | --- | --- | --- | --- |
| | Area ($\text{mm}^2$) | Superior | Middle | Inferior | Area ($\text{mm}^2$) | Superior | Middle | Inferior |
| 35% | 109 | 22% | 29% | 49% | 187 | 17% | 34% | 49% |
| 45% | 112 | 13% | 34% | 53% | 196 | 9% | 44% | 47% |
| 55% | 129 | 38% | 31% | 41% | 230 | 30% | 35% | 35% |
| 65% | 89 | 35% | 21% | 44% | 184 | 24% | 28% | 48% |

**Table 2.**
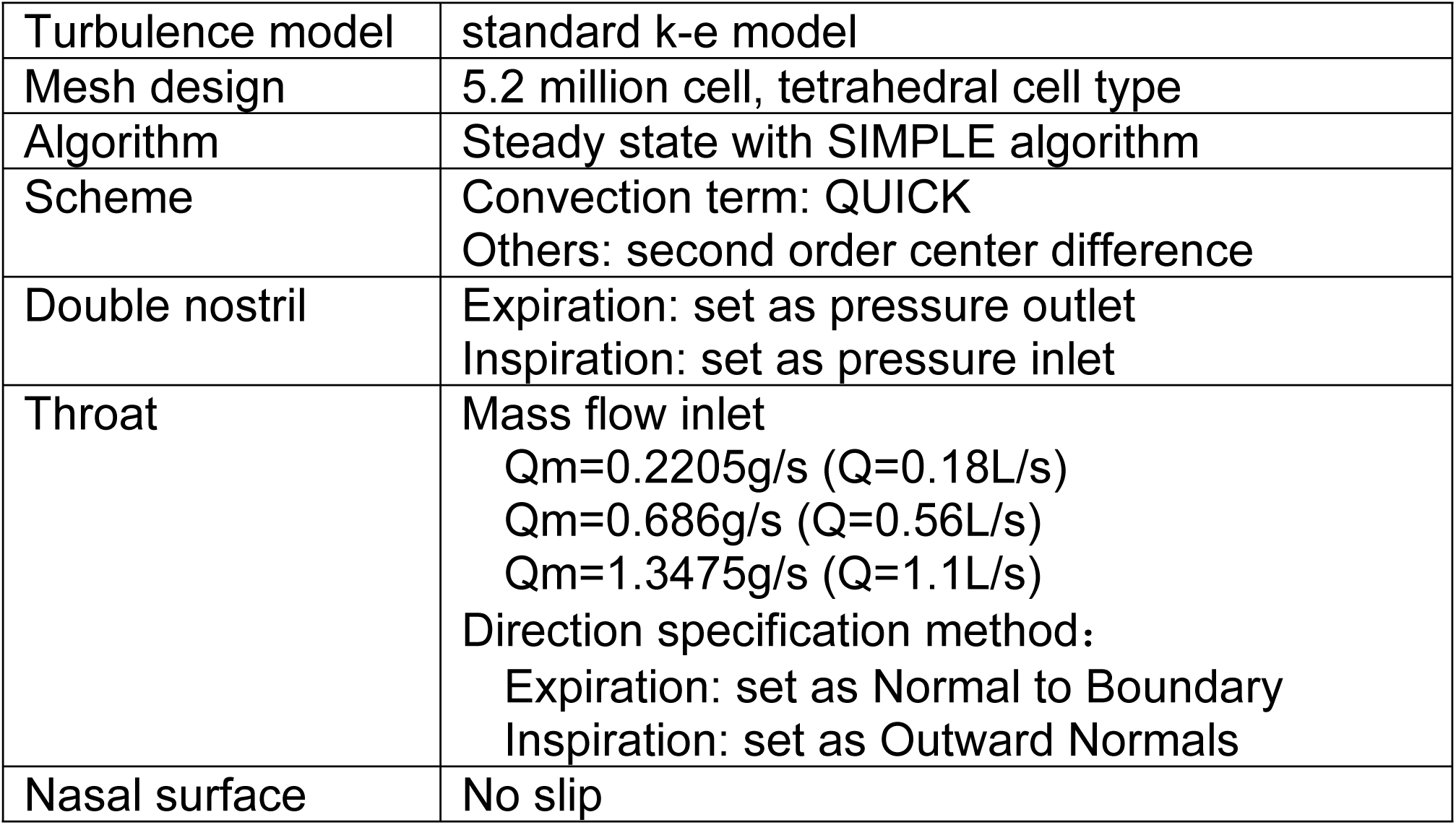
Numerical and boundary conditions

**Fig 5.**
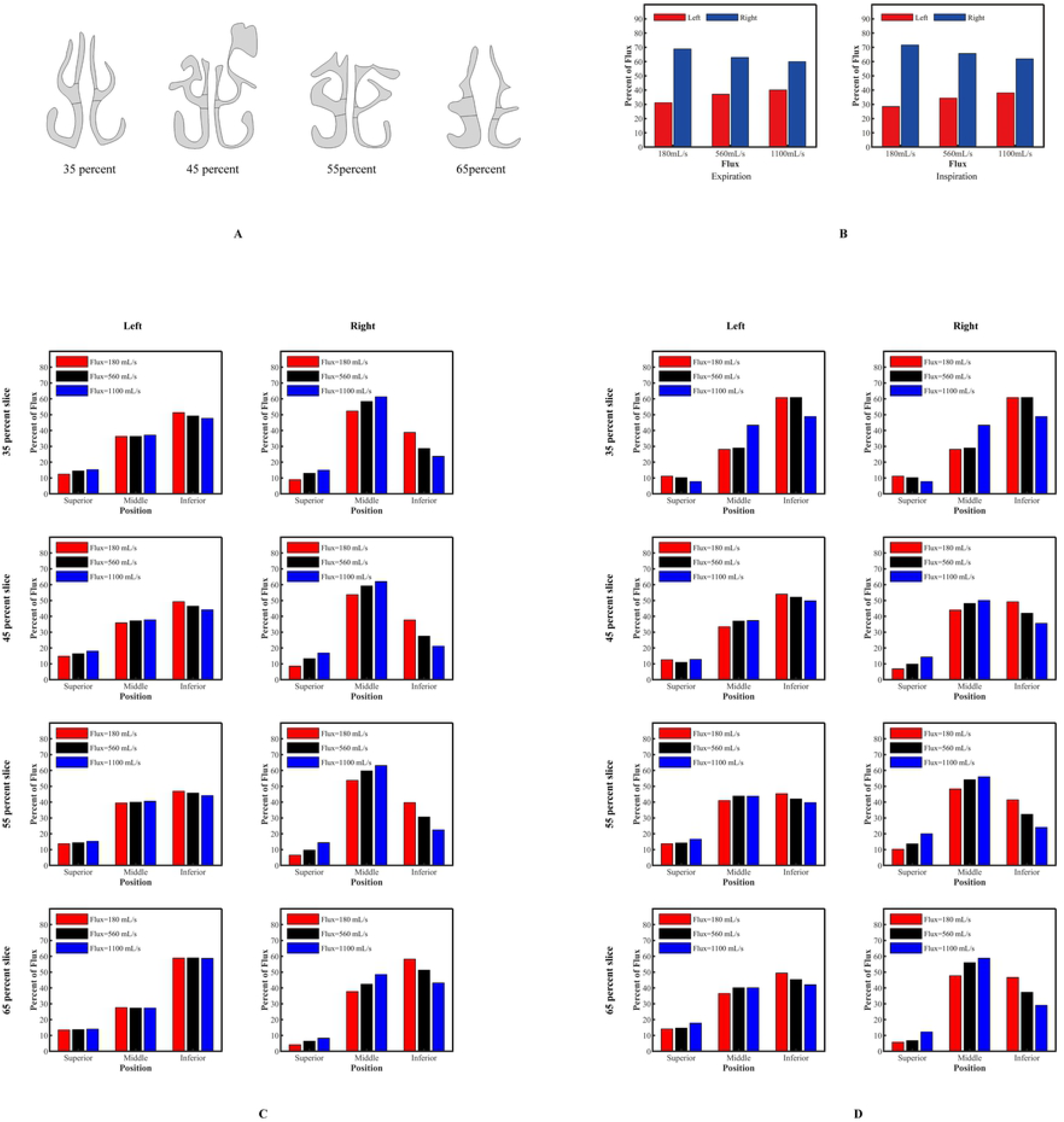
Distributions of the nasal flow. A. The division of the superior, middle, inferior nasal meatus on the coronal section of the nasal cavity at different positions. B. The air distribution between left and right nasal cavity. C. Flow distribution on different sections (along Y-axis direction) at different flow rates (180 ml s^-1^, 560 ml s^-1^ and 1100 ml s^-1^) during inhalation. D. Flow distribution on different sections (along Y-axis direction) at different flow rates during exhalation.

The air distribution between the left and right nasal cavities is shown in Fig 5B. Less flow was distributed to the left nasal cavity than the right one. However, with the increase of the flow, the proportion of flow in the left nasal cavity increased, while that in the right nasal cavity decreased.

Fig 5C shows the distribution of flow at inhalation in the right and left nasal cavities, respectively. The change of flow distribution in the right nasal cavity at different flow rate was more obvious than that in left. The proportion of flow distribution in the right superior and middle nasal meatuses increased with the increase of the flow rate, while the proportion of flow distribution in the right inferior nasal meatus decreased with the increase of the flow rate.

Fig 5D shows the distribution of flow at exhalation. The proportion of flow distribution in the left or right inferior nasal meatus decreased with the increase of the flow. At most sections, the flow proportion of the superior and middle nasal meatuses increased with the increase of the nasal flow. At the 35 percent section, the flow proportion of the left and right superior nasal meatuses decreased with the increase of the nasal flow. At all sections, with the increase of the nasal flow, the flow distribution ratio of the left or right middle nasal meatus increased.

### 6. Stream lines of breath

## Discussion

Diseases of the nasal cavity, especially nasal septum deviation, can affect its airflow. Nasal flow is considered to have close relationship with clinical symptoms, particularly subjective nasal patency[19]. Acoustic rhinometry and rhinomanometry are the commonly used examinations. However, since there is a poor correlation between personal symptoms and examination results, clinical doctors have reservations about these examinations[20]. An increasing number of CFD studies of the flow of the nasal cavity were conducted in order to gain the characteristics of the flow closely related to the symptoms. Nonetheless, few CFD studies have been supported by the experimental data. Pressure is one of the major parameters describing the nasal flow. Accurate pressure is a reliable support for CFD studies, and it is easy to obtain and measure accurately. Therefore, using a normal person as our subject, we examined whether the pressure data we obtained from the CFD study and our experimental data were consistent or not, and further studied the CFD data such as flow velocity and flow volume.

In 2006, Croce et al. [16] created a plastinated model of nasal cavities and maxillary sinuses by using a human anatomical specimen, and studied the pressure of the flow in the nasal cavities and the data of a computer simulation. However, there is no guarantee that the model is in complete agreement with the physiological geometry as the plastinated model, derived from the anatomical specimen, may decrease the thickness of mucous membrane of the turbinates. Besides, the specimen was dehydrated when it was processed. Hahn[4] and Li[15] made the measurement studies of the parameters of the flow using the enlarged models of the nasal cavities. Unlike the models used in the previous studies, the model in our study was obtained from the data of the thin-slice CT scan of a normal human subject in restful state (Fig 1). It was reconstructed meticulously, and maintained almost the complete fine structure of the nasal cavity. Therefore, the model can guarantee the utmost similarity with its physiological geometry. Since the internal structure of the nasal cavities was complicated, we obtained its fine CT image using the thin-slice CT scanning. Most previous CFD studies of the nasal cavities excluded the paranasal sinuses to minimize the reconstruction, meshing, and calculation[12]. In order to get more realistic data, we tried our best to reconstruct the fine internal structure of the nasal cavities, including the maxillary sinus, ethmoid sinus, sphenoid sinus, frontal sinus and its opening based on the CT image in DICOM format. Through threshold segmentation, we obtained the important procedure for the structure of the nasal cavities. The threshold segmentation value is vital to CFD[20, 21]. According to Nakano, the upper threshold value set between −460∼-470 HU has the minimum error[22]. This study chose the upper threshold value of −475 HU. The upper threshold value was manually adjusted near the paranasal sinus opening and in the fine areas of the ethmoid sinus and the bone walls of the sphenoid sinus. So that these areas were not ignored by the automatic threshold segmentation. The precision of the 3-D printing was very high with an error value less than 0.2 mm.

Li et al. used the *k-ε* model, *k-ε* SST model, standard *k-ε* model and LES model to calculate the inferior of the nasal cavities. The comparison with the experimental results indicated that the standard *k-ε* model had a better agreement with the experimental values in velocity profile and turbulence intensity[15]. In light of calculation economy and accuracy, this study used the standard *k-ε* turbulence model.

### Comparison of the calculation results

As the study results show, there is a high agreement between the pressure values at all locations of the nasal cavities in the CFD simulation and the experimental data obtained from the 3-D printed proportional precision model of the nasal cavities.

#### Pressure

This study showed a high agreement of pressure values between the experimental data and the simulation data in most check points especially at the flow rates of 180 ml s^-1^ and 560 ml s^-1^ (Fig 2). Croce et al. [16] studied the inhaling flow volumes at three low flow rates--109, 231, 353 ml s^-1^. The pressure at the flow rate of 180 ml s^-1^ found in this study is consistent with Croce’s results. At the flow rate of 1100 ml s^-1^, there was greater deviation in the experiment data and the simulation data (due to the high deviation at expiration). The possible cause for the deviation was the potential influence on the nasal flow by the walls below the pharynx when the flow rate increased. With the increase of inspiratory or expiratory flow, the pressure drop (or pressure increase) also increased (Fig 2 and 3). This can be analyzed using Bernoulli equation. The increase of flow means the increase of kinetic energy, and the final gas flow rate turns to be 0. Then, this part of kinetic energy was converted into pressure potential energy. The results indicated that the most obvious pressure drop took place in the anterior part of the nasal cavities. The biggest resistance was found in the isthmus nasi, roughly accounting for main part of the total nasal resistance. This is consistent with the previous studies[10, 12]. Solow et al. reported little contribution to the resistance decrease made by the head of the inferior turbinate[2]. Zhao et al. showed that the inferior turbinate contributed far less to the nasal resistance than it was expected[10]. At inspiration, the inferior and middle meatuses had greater pressure drop among the three meatuses whereas that of the superior meatus was relatively gradual. This indicates that the inferior and middle turbinates contribute more to the nasal resistance than the superior turbinate. On the other hand, the gradual pressure drop in the superior meatus might help olfactory perception.

### Flow velocity

There was an obvious difference of flow velocity in space, inspiration, and expiration.

### Spatial difference

In terms of flow rates inside the nasal cavities, the nasal valve area had a higher flow velocity, which is in agreement with the research results of Croce and Zhou[10, 16, 23]. A comparison of the velocity near the three nasal turbinates revealed the following findings: the lowest flow velocity was found near the lateral wall of the inferior turbinate; the medium flow velocity was found near the superior turbinate, and the highest flow velocity was found near the middle turbinate (Fig 4A and B). Zhou et al. [15] believed that the high flow velocity near the middle turbinate correlated with the heat loss and the level of perception of the nasal patency. There was very low flow velocity inside the paranasal sinuses.

### Difference of flow velocity among inspiration and expiration

With the equal quantity of flow for expiration and inspiration, there was a significant difference in the absolute values for the flow velocity at some nasal locations. The flow velocity of inspiration near the middle turbinate and inferior turbinate was significantly larger than that of expiration (Fig 4C). This phenomenon, however, seldom occurred near the nasal septum, suggesting that more flow at inspiration may have heat exchange with the nasal mucosa. This facilitated the cooling of the nasal mucosa and the sensation of the air temperature by the trigeminal thermoreceptors in the nasal mucosa[24], which might be related to the subjective feeling of the nasal flow obstruction. The inhaled flow of the superior meatus was also much greater than the exhaled flow, which might henlp the olfactory area to get more odor molecules during inspiration than expiration.

### Airflow distribution

Borojeni et al. reported that for human individuals the middle meatus flow and inferior meatus flow were significantly correlated with total ipsilateral flow whereas the superior meatus flow did not correlate significantly with ipsilateral airflow[12]. Casey’s study reported that the middle meatus flow in healthy individuals was significantly higher than that in people with septal deviations (narrow side). Additionally, subjective nasal patency did not correlate with the inferior meatus flow and the superior meatus flow, but was correlated with the middle meatus flow[19]. Our study (Fig 5) found that as the breathing flow of the bilateral nasal cavities increased, the flow ratio of the inferior meatus to the ipsilateral nasal cavity tended to decrease (only 65% of the left-side cross section did not change). On the other hand, the flow ratio of the middle meatus tended to increase significantly. It means that more air flow was distributed to middle meatus if breathed deeply. The flow ratio of the superior meatus had different change patterns at expiration and inspiration. At inspiration, like that in the middle meatus, the flow ratio at all cross sections of the superior meatus increased. At expiration, with the increase of the flow volume, the flow ratio in the posterior slices of the superior meatus tended to increase whereas the flow ratio in the anterior slices of the meatus tended to decrease. In other words, the overall increase of the flow volume in the nasal cavities was not equally distributed among the superior, middle, and inferior meatuses. The middle meatus had the most obvious increase, which might help it better perform the functions of cooling, sensing nasal air flowing, and ventilating in the paranasal sinuses.

In general, the flow ratio in the superior meatus was around 10%. This is in agreement with a previous study[2]. The increase of the flow ratio at inspiration might be conducive to the sensation of smell and ventilation in the sphenoid sinus and the posterior ethmoid sinus opened in the back part of the superior meatus. The increase of the flow ratio in the posterior part of the superior meatus at expiration might facilitate the ventilation in the sphenoid sinus and the posterior ethmoid sinus. The decrease in flow ratio in the anterior part of the superior meatus (close to the olfactory regions) at expiration could be related to the fact that the sensation of breathing-out3 smell was not a major function of the nasal cavities. The characteristics of the airflow distribution could be attributed to the formation of the nasal cavities during evolution.

### Streamlines

Li et al. observed significant turbulence in the anterior dorsal nasal cavity and the pharynx channel when the flow rate increased to 560 ml s^-1^, and considerable turbulence flow around the middle and inferior turbinate regions when the flow rate was 1100 ml s^-1^ [15]. Our study found that the area and scale of the turbulence regions increased as the flow volume increased (Fig 6). We also found significant inspiratory flow, named as “inspiratory jet” by Hildebrandt’s study[2], from the nasal valve to the middle and inferior meatus. Tan et al. reported that there was less turbulent flow in the posterior part of the nasal cavities in Chinese people compared with that in westerners[23]. Our study found obvious formation of turbulence in the nasopharynx area, which has plenty of lymphatic tissues and full interactions with the nasal flow. The formation of the turbulence there could be attributed to the immune response and inflammation in the area. At expiration, separate turbulence regions were created by the flow close to the soft palate. As a result, a pressure difference near the walls of the soft palate and the major flow area was formed. The pressure difference made the soft palate vibrate, causing snoring.

**Fig 6.**
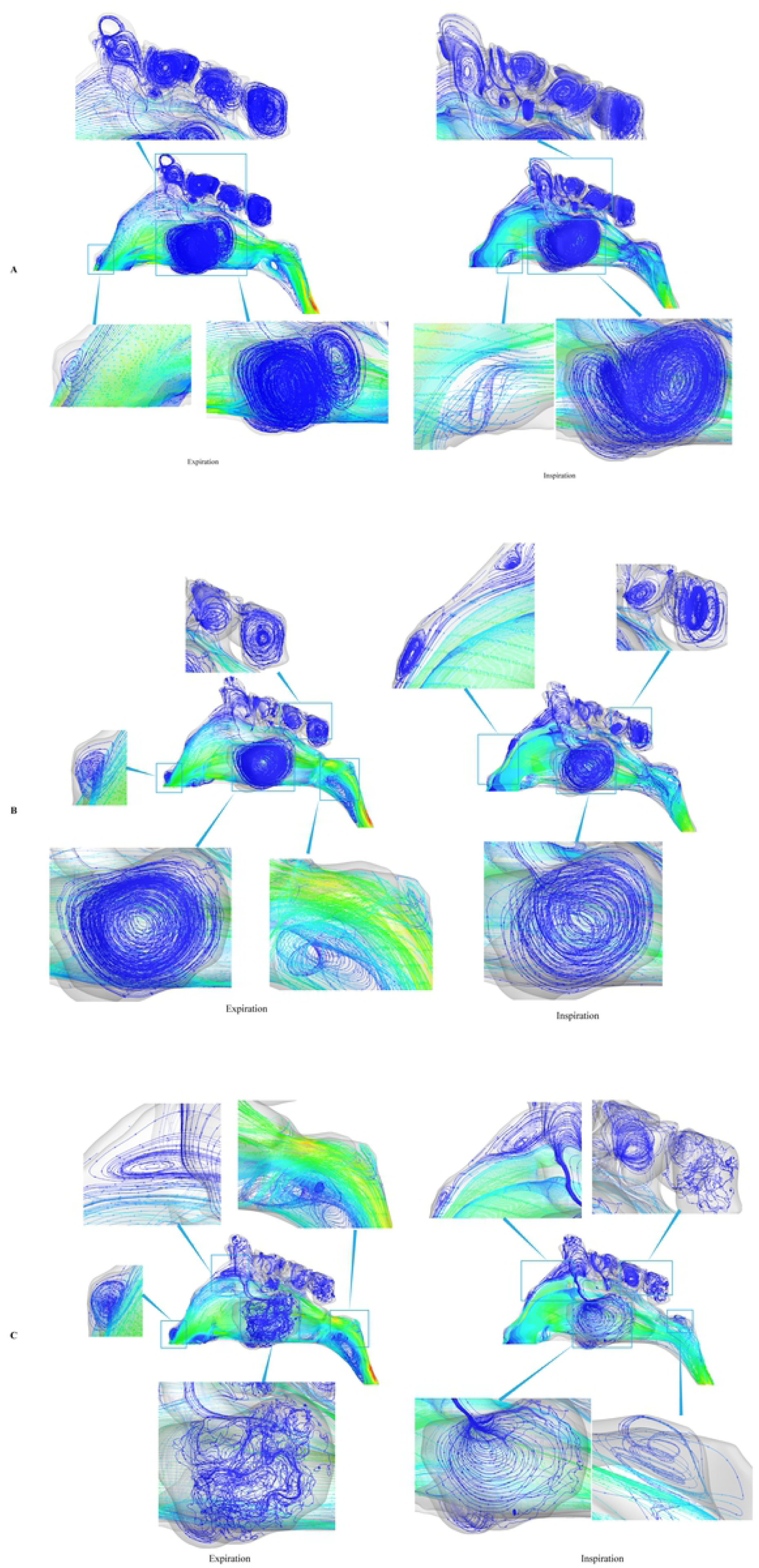
Stream lines of nasal flow. A. Stream lines in nasal cavity of flow at 180 ml s^-1^. B. Stream lines in nasal cavity of flow at 560 ml s^-1^. C. Stream lines in nasal cavity of flow at 1100 ml s^-1^. Fig 6 shows the stream lines of the nasal flow at three different flow rates. Note that at all these flow rates, vortex formed in the nasal cavity, especially in the nasal valve, the olfactory regions, and the nasopharynx area. At the flow rate of 180 ml s^-1^, the flow field was relatively stable. With the increase of the flow rate, the flow lines became disordered, meaning the flow would be violent and turbulent.

Li et al. found an anterior airflow vortex at all nasal rates, which could cause further mixing of the flow and intensifying its unsteadiness in the region[15]. Wen et al. also found vortex formations posterior to the nasal valve and olfactory regions at the flow rates of 7.5 L min^-1^. and 15 L min^-1^ [25]. In our study, the flow in the olfactory regions was relatively slow, which might facilitate the sensation of odorants. Turbulence was observed in the anterior part of the nasal cavities close to the olfactory regions at the flow rate of 560 ml s^-1^ for the inspiratory air (Fig 6B). It was possible that the turbulence was captured by the mucosa receptors in the olfactory regions where they helped the odorants stay completely. One study reported that inspiration at the flow rate around 560 ml s^-1^ could facilitate the sensation of smell[15]. Thus, the inspiratory flow rate of 560 ml s^-1^ was also called medium sniffing breathing condition[15]. No turbulence was observed in the anterior part of the nasal cavities at the flow rates of 180 ml s^-1^ and 1100 ml s^-1^ in this study. As for the results of the streamlines, the enlarged model of the nasal cavity used by Li et al. [15], just like the one used by Hahn in 1993[4], could be the reason for the difference between our results and Li et al.’s results. As for the function of the turbulence in sensing smell, Zhao et al reported that the olfactory mucosal uptake of smell created by the mixed turbulence at high flow rates could be negligible as sniff duration was more important than sniff strength[26]. Therefore, the relationship between turbulence and olfactory detection needs to be further studied.

The limitations of the study are: First of all, since our model was 3-D printed using resin as its material, it was harder than the real nasal mucosa in terms of texture. Changing the resin model into a silicon model can make the edges more similar to the texture of the nasal mucosa, thus making the measurements closer to reality. Second, the pressure of the nasal cavity was measured at steady nasal flow rates. However, the real breathing rates in the nasal cavity are cyclical, with periodical interactions among nasal flows. So, it probably will be more accurate to measure the pressure at the cyclical breathing rates.

In conclusion, the computational methods and boundary conditions used in this study produced a good agreement between the CFD simulation results and the experimental results. The fine and precise reconstruction of the nasal cavities helped the study of the influence of the internal nasal structure on the nasal flow. The method used in this study to measure the pressure in the nasal cavities can be used in experimental measurements of the partial resistance at almost all locations of the nasal cavity. Unlike the rhinomanometry, which can only measure a single whole resistance of the nasal cavity, our method can also be used in the validation of CFD studies of the nasal cavity. With proper modification, it can be applied to the clinical practice for nasal resistance, giving more help for the design of the operation plan.

## Materials and methods

### Ethics Statement

This study and the use of human clinical digitals were approved by the Institutional Research Ethics Committee of the Second Military Medical University, Shanghai, China. Investigations were conducted according to the principles expressed in the Declaration of Helsinki. The patient, without history of nasal diseases, provided written informed consent.

### CT scanning

One normal subject was chosen (female, aged 39). The subject was in restful state (rested for 15 minutes). The CT scan of the subject’s head was taken in supine position, with the slice thickness of 0.625mm.

### Creation of the model for the nasal cavities and the paranasal sinuses

The CT scan file in DCOM format was imported into the MIMICS software (19.0). The segmentation threshold value was set at −1024∼-475 HU. The file was manually cropped. The model of the nasal cavities, the paranasal sinuses, and the pharynx was obtained and exported as a model in IGES format. The accuracy of the model was checked by an experienced engineer in nasal anatomy and a certified ENT doctor.

### Grid for the model of the nasal cavities and paranasal sinuses

Pointwise was used to generate a grid to calculate the internal airflow of the nasal cavities. As the model was for the complete nasal cavities, which have a complex structure, a tetrahedral unstructured grid was used as the grid type to ensure its efficiency. Finely generated, the mesh well reflected the internal features of the nasal cavities.

### Computer simulation of the model of the nasal cavities and paranasal sinuses

The grid was imported into Ansys Fluent to find the solutions to the airflow of the nasal cavities. The flow inside the nasal cavities was calculated in its steady state. *K-ε* model was used for the simulation of turbulent flow. The CFD methods were used for studying the airflow of the nasal cavities at three flow rates of 180 ml s^-1^, 560 ml s^-1^,and 1100 ml s^-1^, corresponding to restful breathing, medium sniffing, and strong sniffing[15].

Table 2 is the detailed parameter setting from Fluent. In exhaling state, the boundary conditions for the nostrils were set as “pressure outlet.” In inhaling state, the boundary conditions were set as “pressure inlet.”

### Experimental measurement of the pressure in the nasal cavities

Holes were punched inside the virtual model of the nasal cavities and the paranasal sinuses in IGES format. The channels of the holes connected the point we studied inside the nasal cavities and the pressure check points at the surface of the model. The pressure check points are shown exactly in Fig 1. The model was printed by a high precision 3-d printer (photosensitive resin, DSM Somos 8000, tolerance of dimension 0.2mm, UnionTechmaterial, machine model). An air compressor (DET-1500, Daertuo) and a vacuum (VP-1500, Daertuo) were used to make steady air flow in nasal cavity model.

## Acknowledgments

Thanks for the image collection work of Dr. He Xu and Dr. Hua Li.

## Uncategorized References

1. Quadrio M, Pipolo C, Corti S, Lenzi R, Messina F, Pesci C, et al. Review of computational fluid dynamics in the assessment of nasal air flow and analysis of its limitations. Eur Arch Otorhinolaryngol. 2014;271(9):2349-54. Epub 2013/10/09. doi: 10.1007/s00405-013-2742-3. PubMed PMID: 24100883.

2. Hildebrandt T, Goubergrits L, Heppt WJ, Bessler S, Zachow S. Evaluation of the intranasal flow field through computational fluid dynamics. Facial plastic surgery: FPS. 2013;29(2):93-8. Epub 2013/04/09. doi: 10.1055/s-0033-1341591. PubMed PMID: 23564240.

3. Solow B, Sandham A. Nasal airflow characteristics in a normal sample. Eur J Orthod. 1991;13(1):1-6. Epub 1991/02/01. doi: 10.1093/ejo/13.1.1. PubMed PMID: 2032561.

4. Hahn I, Scherer PW, Mozell MM. Velocity profiles measured for airflow through a large-scale model of the human nasal cavity. J Appl Physiol (1985). 1993;75(5):2273-87. Epub 1993/11/01. doi: 10.1152/jappl.1993.75.5.2273. PubMed PMID: 8307887.

5. Churchill SE, Shackelford LL, Georgi JN, Black MT. Morphological variation and airflow dynamics in the human nose. Am J Hum Biol. 2004;16(6):625-38. Epub 2004/10/21. doi: 10.1002/ajhb.20074. PubMed PMID: 15495233.

6. Weinhold I, Mlynski G. Numerical simulation of airflow in the human nose. Eur Arch Otorhinolaryngol. 2004;261(8):452-5. Epub 2003/12/04. doi: 10.1007/s00405-003-0675-y. PubMed PMID: 14652769.

7. Muller-Wittig W, Mlynsji G, Weinhold I, Bockholt U, Voss G. Nasal airflow diagnosis--comparison of experimental studies and computer simulations. Stud Health Technol Inform. 2002;85(undefined):311-7. Epub 2004/10/02. doi: 10.3233/978-1-60750-929-5-311. PubMed PMID: 15458107.

8. 1. 8. Leong SC, Chen XB, Lee HP, Wang DY. A review of the implications of computational fluid dynamic studies on nasal airflow and physiology. Rhinology. 2010;48(2):139-45. Epub 2010/05/27. doi: 10.4193/Rhin09.133. PubMed PMID: 20502749.

9. Bailie N, Hanna B, Watterson J, Gallagher G. An overview of numerical modelling of nasal airflow. Rhinology. 2006;44(1):53-7. Epub 2006/03/23. PubMed PMID: 16550951.

10. Zhao K, Jiang J. What is normal nasal airflow? A computational study of 22 healthy adults. Int Forum Allergy Rhinol. 2014;4(6):435-46. Epub 2014/03/26. doi: 10.1002/alr.21319. PubMed PMID: 24664528; PubMed Central PMCID: PMCPMC4144275.

11. Lu J, Han D, Zhang L. Accuracy evaluation of a numerical simulation model of nasal airflow. Acta oto-laryngologica. 2014;134(5):513-9. Epub 2014/04/08. doi: 10.3109/00016489.2013.863430. PubMed PMID: 24702230.

12. Borojeni AAT, Garcia GJM, Moghaddam MG, Frank-Ito DO, Kimbell JS, Laud PW, et al. Normative ranges of nasal airflow variables in healthy adults. Int J Comput Assist Radiol Surg. 2020;15(1):87-98. Epub 2019/07/04. doi: 10.1007/s11548-019-02023-y. PubMed PMID: 31267334; PubMed Central PMCID: PMCPMC6939154.

13. Sommer F, Hoffmann TK, Mlynski G, Reichert M, Grossi AS, Kroger R, et al. [Three-dimensional analysis of nasal physiology: Representation by means of computational fluid dynamics]. HNO. 2018;66(4):280-9. Epub 2017/12/10. doi: 10.1007/s00106-017-0443-8. PubMed PMID: 29222682.

14. Mosges R, Buchner B, Kleiner M, Freitas R, Horschler I, Schroder W. Computational fluid dynamics analysis of nasal flow. B-ent. 2010;6(3):161-5. Epub 2010/11/26. PubMed PMID: 21090156.

15. Li C, Jiang J, Dong H, Zhao K. Computational modeling and validation of human nasal airflow under various breathing conditions. J Biomech. 2017;64:59-68. Epub 2017/09/13. doi: 10.1016/j.jbiomech.2017.08.031. PubMed PMID: 28893392; PubMed Central PMCID: PMCPMC5694356.

16. Croce C, Fodil R, Durand M, Sbirlea-Apiou G, Caillibotte G, Papon JF, et al. In vitro experiments and numerical simulations of airflow in realistic nasal airway geometry. Ann Biomed Eng. 2006;34(6):997-1007. Epub 2006/06/20. doi: 10.1007/s10439-006-9094-8. PubMed PMID: 16783655.

17. Mylavarapu G, Murugappan S, Mihaescu M, Kalra M, Khosla S, Gutmark E. Validation of computational fluid dynamics methodology used for human upper airway flow simulations. J Biomech. 2009;42(10):1553-9. Epub 2009/06/09. doi: 10.1016/j.jbiomech.2009.03.035. PubMed PMID: 19501360.

18. Hooper RG, editor Forced inspiratory nasal flow-volume curves: a simple test of nasal airflow. Mayo Clin Proc; 2001: Elsevier.

19. Casey KP, Borojeni AA, Koenig LJ, Rhee JS, Garcia GJ. Correlation between Subjective Nasal Patency and Intranasal Airflow Distribution. Otolaryngol Head Neck Surg. 2017;156(4):741-50. Epub 2017/02/01. doi: 10.1177/0194599816687751. PubMed PMID: 28139171; PubMed Central PMCID: PMCPMC6062004.

20. Quadrio M, Pipolo C, Corti S, Messina F, Pesci C, Saibene AM, et al. Effects of CT resolution and radiodensity threshold on the CFD evaluation of nasal airflow. Medical & biological engineering & computing. 2016;54(2-3):411-9. Epub 2015/06/11. doi: 10.1007/s11517-015-1325-4. PubMed PMID: 26059996.

21. Cherobin GB, Voegels RL, Gebrim E, Garcia GJM. Sensitivity of nasal airflow variables computed via computational fluid dynamics to the computed tomography segmentation threshold. PLoS One. 2018;13(11):e0207178. Epub 2018/11/18. doi: 10.1371/journal.pone.0207178. PubMed PMID: 30444909; PubMed Central PMCID: PMCPMC6239298.

22. Nakano H, Mishima K, Ueda Y, Matsushita A, Suga H, Miyawaki Y, et al. A new method for determining the optimal CT threshold for extracting the upper airway. Dentomaxillofac Radiol. 2013;42(3):26397438. Epub 2012/07/31. doi: 10.1259/dmfr/26397438. PubMed PMID: 22842640; PubMed Central PMCID: PMCPMC3667543.

23. Tan J, Han D, Wang J, Liu T, Wang T, Zang H, et al. Numerical simulation of normal nasal cavity airflow in Chinese adult: a computational flow dynamics model. Eur Arch Otorhinolaryngol. 2012;269(3):881-9. Epub 2011/09/23. doi: 10.1007/s00405-011-1771-z. PubMed PMID: 21938528.

24. Sozansky J, Houser SM. The physiological mechanism for sensing nasal airflow: a literature review. Int Forum Allergy Rhinol. 2014;4(10):834-8. Epub 2014/08/01. doi: 10.1002/alr.21368. PubMed PMID: 25079504.

25. Wen J, Inthavong K, Tu J, Wang S. Numerical simulations for detailed airflow dynamics in a human nasal cavity. Respir Physiol Neurobiol. 2008;161(2):125-35. Epub 2008/04/02. doi: 10.1016/j.resp.2008.01.012. PubMed PMID: 18378196.

26. Zhao K, Dalton P, Yang GC, Scherer PW. Numerical modeling of turbulent and laminar airflow and odorant transport during sniffing in the human and rat nose. Chem Senses. 2006;31(2):107-18. Epub 2005/12/16. doi: 10.1093/chemse/bjj008. PubMed PMID: 16354744.

